# Identification of functional rare coding variants in IGF-1 gene in humans with exceptional longevity

**DOI:** 10.1101/2024.10.11.617885

**Authors:** Amanat Ali, Zhengdong Zhang, Tina Gao, Sandra Aleksic, Evripidis Gavathiotis, Nir Barzilai, Sofiya Milman

## Abstract

Diminished signaling via insulin/insulin-like growth factor-1 (IGF-1) axis is associated with longevity in different model organisms. IGF-1 gene is highly conserved across species, with only few evolutionary changes identified in it. Despite its potential role in regulating life span, no coding variants in IGF-1 have been reported in human longevity cohorts to date. This study investigated the whole exome sequencing data from 2,487 individuals in a cohort of Ashkenazi Jewish centenarians, their offspring, and controls without familial longevity to identify functional IGF-1 coding variants. We identified two likely functional coding variants *IGF-1*:p.Ile91Leu and *IGF-1*:p.Ala118Thr in our longevity cohort. Notably, a centenarian specific novel variant *IGF-1:p*.Ile91Leu was located at the binding interface of IGF-1 – IGF-1R, whereas *IGF-1*:p.Ala118Thr was significantly associated with lower circulating levels of IGF-1. We performed extended all-atom molecular dynamics simulations to evaluate the impact of Ile91Leu on stability, binding dynamics and energetics of IGF-1 bound to IGF-1R. The *IGF-1*:p.Ile91Leu formed less stable interactions with IGF-1R’s critical binding pocket residues and demonstrated lower binding affinity at the extracellular binding site compared to wild-type IGF-1. Our findings suggest that *IGF-1*:p.Ile91Leu and *IGF-1*:p.Ala118Thr variants attenuate IGF-1R activity by impairing IGF-1 binding and diminishing the circulatory levels of IGF-1, respectively. Consequently, diminished IGF-1 signaling resulting from these variants may contribute to exceptional longevity in humans.

## Introduction

Diminished signaling via insulin/insulin-like growth factor 1 (IGF-1) axis has been associated with increased lifespan in various model organisms^1-4^. However, the role of insulin/IGF-1 axis in human aging has not been confirmed. Previously, two rare heterozygous coding variants in *IGF-1R*, a gene that encodes the IGF-1 receptor (IGF1-R), were found to be enriched among Ashkenazi Jewish individuals with exceptional longevity and were demonstrated to result in diminished activity of the IGF-1R^5^. However, IGF-1 is a highly conserved gene and the few missense mutations identified in *IGF-1* to date have been associated with growth failure and developmental abnormalities^6-8^. To our knowledge, no coding variants in the *IGF-1* have been associated with longevity in humans.

IGF-1 mediated downstream signaling is dependent on stable binding of IGF-1 with its receptor IGF-1R. Studies have shown that coding variants located at the interface of IGF-1 - IGF-1R attenuated the binding activity of IGF-1^9,10^. The IGF-1R is a functional dimer. Each protomer of IGF-1R includes the L1 (leucine-rich repeat domain 1), CR (cysteine-rich domain), L2 (leucine-rich repeat domain 2), FnIII-1, -2, -3 (fibronectin type III domains), transmembrane (TM), a ∼30 amino acid juxtamembrane region, and kinase domains. Two of these protomers are connected by numerous disulfide bonds, creating a stable, covalent dimer^11^. For clarity, the domains in protomers 1 and 2 are indicated as L1 – FnIII-3, and L1′ – FnIII-3′ (indicated by prime), respectively, throughout the manuscript. Ligand binding to the extracellular domains (ECDs) of IGF-1R triggers receptor kinase activation that results in the phosphorylation of numerous substrates and the initiation of distinct signaling pathways^12^. Members of the insulin receptor (IR) family stand out among receptor tyrosine kinases (RTKs) by forming dimers composed of αβ subunits. Each αβ dimer possesses two ligand-binding sites. Each site comprises two distinct partial sites referred to as site 1 and site 2. Site 1 is formed by residues on L1 from one subunit and residues on the αCT′ helix of the other subunit (in model organisms) or αCT helix of the same subunit (in human), while site 2 consists of residues on Fn1′ and Fn2′^13-16^. In addition to the well-characterized primary IGF-1-binding site^11,17-19^ that comprises the L1 domain and α-CT, a secondary sub-site has recently been observed in the active IGF-1R dimer^11^. However, there is limited understanding regarding the stability and significance of polar and hydrophobic contacts established between wild-type and mutant IGF-1 and IGF-1R. Furthermore, regulation of IGF-1 signaling involves alternative splicing that results in different IGF-1 precursors which vary in the structure of their carboxy-terminal extension peptides (E-peptides) and the length of their amino-terminal signal peptides^20^. Different splicing variants and synonymous variants have been shown to alter the expression, function, and processing of mature IGF-1^21-23^.

Multiple three-dimensional atomic-level structures of IGF-1 bound to IGF-1R have been successfully elucidated^11,17,19,24^. These resolved structures offer profound insights into macromolecular structure and intermolecular interactions. Yet, molecular recognition and binding involve dynamic processes. Molecular dynamic (MD) simulations often serve as a complement to conventional structural studies, allowing for the examination of these processes at atomic-level^25,26^. These simulations offer insights into the stability of macromolecular complexes, the flexibility of interacting subunits, and the interactions among residues at the binding interface.

In this study, we investigated the impact of longevity-associated *IGF-1* coding variants identified in individuals with exceptional longevity. We associated *IGF*-1 variants with serum IGF-1 levels and determined the effect of interfacial variant *IGF-1*:p.Ile91Leu on stability, binding dynamics, and energetics of IGF-1 bound to IGF-1R by performing extended MD simulations. The main aim of this study was to uncover both commonalities and disparities in the dynamic interactions between wild-type and longevity-associated mutant IGF-1, as well as pinpoint residues that may play a pivotal role in maintaining the integrity of this interface in the presence of studied variants.

## Results

### Identification of functional variants in IGF-1 gene

We studied the whole exome sequencing data from 2,487 individuals in a cohort of Ashkenazi Jewish centenarians, offspring of centenarians, and offspring of parents without familial longevity to identify all coding variants in the *IGF-1* gene. The characteristics of the longevity cohort are summarized in **Supplementary Table S1**. Only two coding variants were identified in our cohort and both had minor allele frequency (MAF) ≤ 0.01 **(Figure 1A)**. The functional nature of the coding variants was defined using combined annotation dependent depletion (CADD) score^27^. Variants with CADD score ≥ 20 were considered functional. Both identified variants had CADD scores ≥ 20 and were present in a heterozygous state. A novel variant *IGF-1:p*.Ile91Leu was found in two female centenarians, whereas the *IGF-1*:p.Ala118Thr variant was found in two centenarians as well as in four offspring and three control individuals. The latter variant was classified by ClinVar as a variant of unknown significance (VUS) that was previously noted in probands with growth delay due to IGF-1 deficiency. IGF-1 gene is conserved among mammals^28^. IGF-1 protein sequences of different mammals obtained from UniProt and ClustalW^29^ was employed for multiple sequence alignment (MSA) to assess the conservation of Ile91 and Ala118 residues. Both Ile91 and Ala118 are highly conserved (**Supplementary Figure S1**). Thus, substitutions at these positions could potentially have detrimental effects on the protein’s structure and function. In our cohort, carriers of *IGF-1*:p.Ile91Leu variants had insignificantly lower maximal reported height compared to non-carriers, adjusted for sex, (158.7 ± 1.8 cm vs. 163.4 ± 9.0 cm, respectively, p=0.36), while the maximal reported height was comparable between *IGF-1*:p.Ala118Thr carriers and non-carriers (166.1 ± 7.1 cm vs. 166.7 ± 9.9 cm, respectively, p=0.84). The serum IGF-1 levels of *IGF-1:p*.Ile91Leu carriers were not measured. Interestingly, the carriers of *IGF-1*:p.Ala118Thr had significantly lower levels of IGF-1 compared to non-carriers (**Figure 1B**).

**Figure 1.**
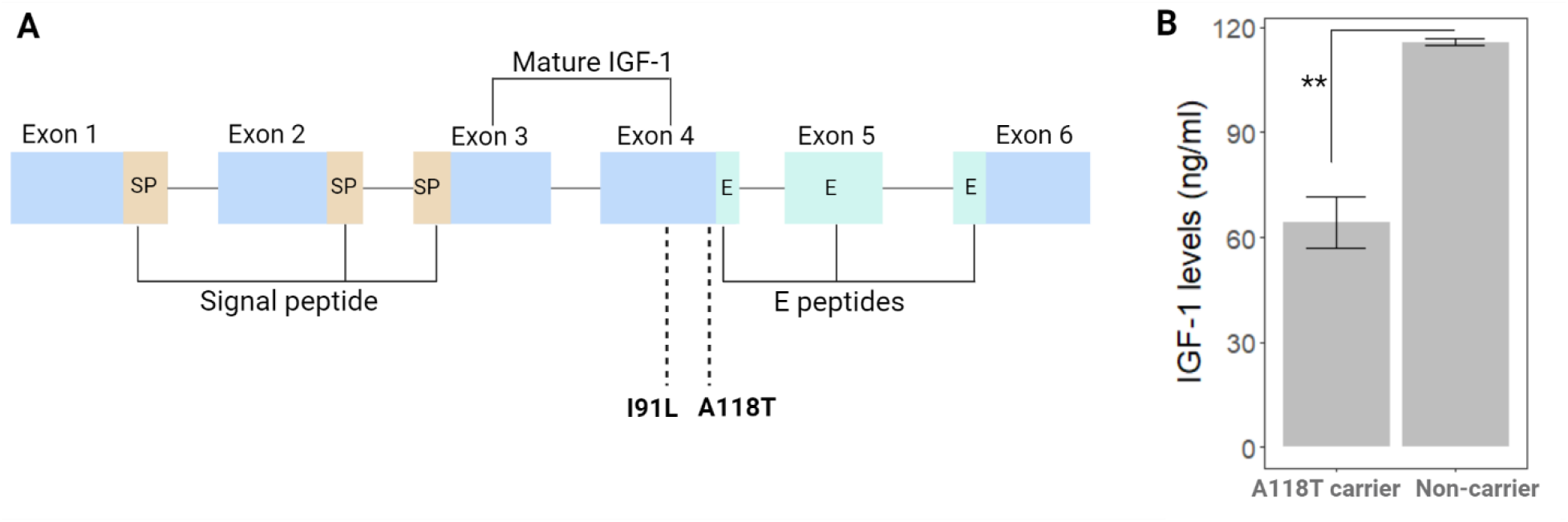
**A)** The structure of IGF-1 gene. Our longevity cohort carries two missense variants in exon 4. Brown, light cyan and blue colors represent the signal peptide (SP), E peptide and protein-coding regions, respectively. A118T variant is located at the boundary of Exon 4 and the N-terminal sequence of the E-peptides, which is a cleavage site for the release of mature IGF-1. B) Association of IGF-1:p.Ala118Thr with serum IGF-1 levels. Results are plotted as mean ± SD. The statistical model was adjusted with baseline age and sex. * * p<0.01.

### Impact of mutants on IGF-1 – IGF-1R bound structure

In order to ascertain the role of missense variants in perturbing the IGF-1 - IGF-1R architecture and the binding of IGF-1 to IGF-1R, we investigated the impact of *IGF-1* gene variants on IGF-1R structure and function. *IGF-1:p*.Ile91Leu variant was located at the binding interface of IGF-1 – IGF-1R **(Figures 2A-2C)**. At the static structure level, neither isoleucine nor leucine established contacts with the adjacent residues of IGF-1R Ile91Leu was tracked during MD simulations to observe its binding potential with neighboring residues of IGF-1R and IGF-1. In the wild-type runs, Ile91 formed more sustained interaction with the critical binding pocket residue Phe731of IGF-1R when compared to mutant runs (Leu91). In contrast, Leu91 exhibited consistent interactions with Glu94 and Cys95 of IGF-1, unlike the wild-type Ile91 **(Figures 2B-2D)**. This suggested that Leu91 variant was more engaged in forming interactions with neighboring residues of IGF-1 and was less readily available for forming interaction with IGF-1R compared to wild-type Ile91 residue. *IGF-1*:p.Ala118Thr, on the other hand, was found at the C-terminal end of the IGF-1 molecule, a region that is not involved in IGF-1R binding. As expected, Ala118Thr did not establish contact with residues of IGF-1R **(Figures 2E and 2F)**.

**Figure 2.**
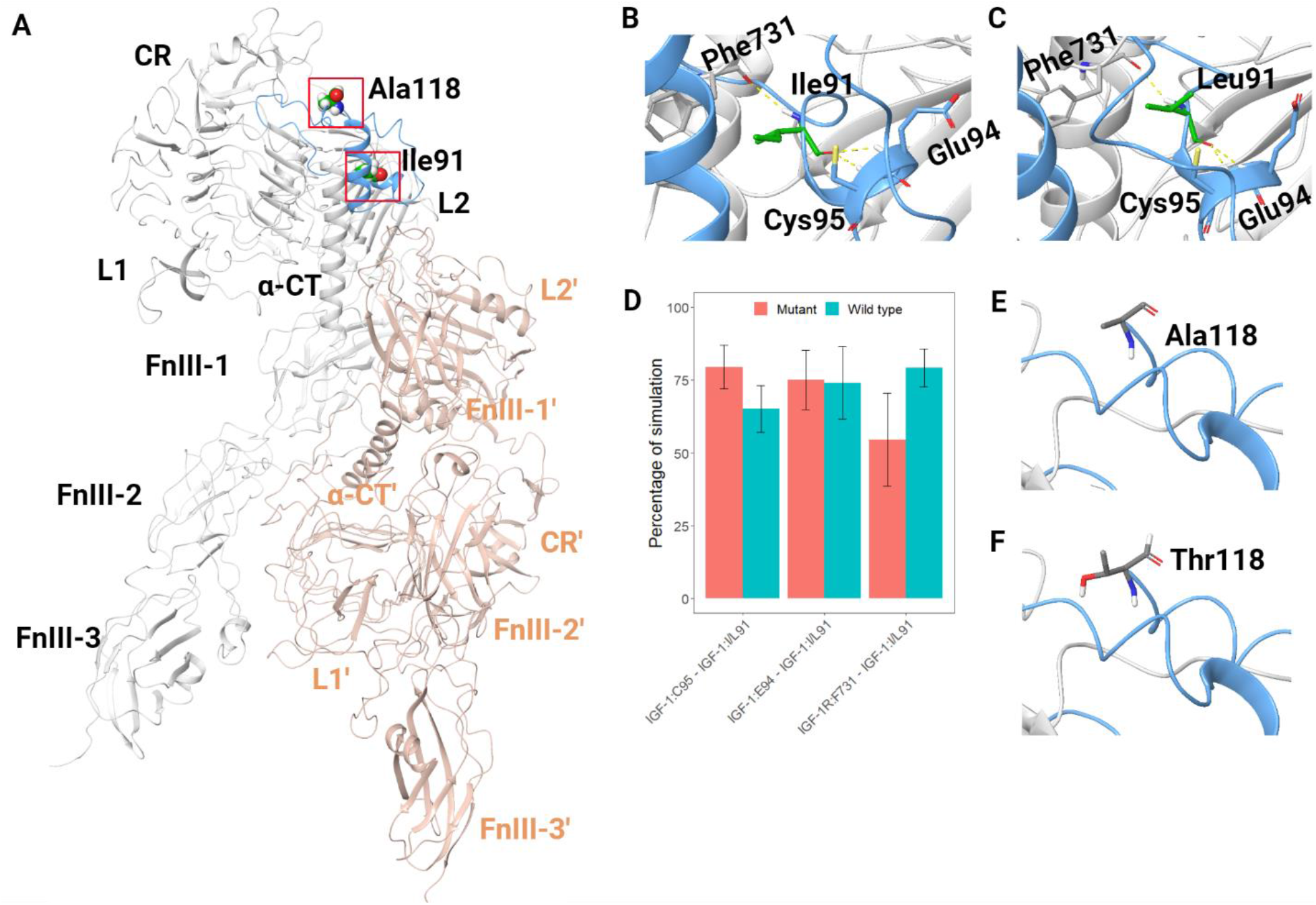
Three-dimensional structure of human IGF-1R bound to IGF-1. The red-boxed regions in (A) are magnified in the successive images (B, C, D and E). The IGF-1R dimer is depicted with two chains, displayed in gray and light pink, while IGF-1 is illustrated with a blue cartoon. A) Structure of IGF-1R bound with IGF-1; B) Wild-type Ile91; C) Mutant Leu91; D) The percentage of simulation time during which Ile91Leu maintained contacts with neighboring residues of IGF-1R and IGF-1.E) Wild-type Ala118; F) Mutant Thr118. Hydrogen bonds are represented with yellow dotted lines.

### MD simulations of wild-type and mutant IGF-1 – IGF-1R complexes

Given that the *IGF-1*.Ile91Leu variant was located at the binding interface of IGF-1 and IGF-1R, we conducted extended MD simulations to gain mechanistic insights into how this variant may affect the binding architecture of IGF-1 with IGF-1R. *IGF-1*:p.Ala118Thr was excluded from MD simulation because it was situated at the boundary of the mature IGF-1 molecule and was not demonstrated to be important for IGF-1R binding. MD simulations of wild-type and *IGF-1*.Ile91Leu (referred to as mutant IGF-1 here onwards) complexes of IGF-1 – IGF-1R were performed in triplicates, each of 500 ns duration, to avoid bias in results often caused by a single simulation run. Runs of the wild-type and mutant simulations were extended to 500 ns to ensure that simulations remained stable and the interactions were faithfully retained for longer duration. The root mean square deviation (RMSD) results from the three runs per simulated system were averaged, and the mean evolution for each system, along with the standard deviations, are shown in **Figure 3**. The individual contributions of the different runs for each system are provided in **Supplementary Figure S2**. The overall structural integrity of all simulations of wild-type and mutant complexes remained stable with a Cα RMSD from the initial structure that was less than 11 Å **(Figure 3A)**. Both wild-type and mutant complexes reached equilibrium after a few nanoseconds of simulation and remained stable throughout the course of simulations. IGF-1R is a dimeric macromolecule and is expected to produce slightly higher RMSD than monomeric structures. Nevertheless, all the runs eventually reached convergence by the end of the simulations. Simulations of dimeric proteins as well as mutated proteins have shown similarly elevated RMSD values^30,31^. The secondary structure composition and compactness of the IGF-1R and IGF-1 protein structures, as represented by the radius of gyration, remained preserved throughout the simulations **(Supplementary Table S2 and Table S3)**.

**Figure 3.**
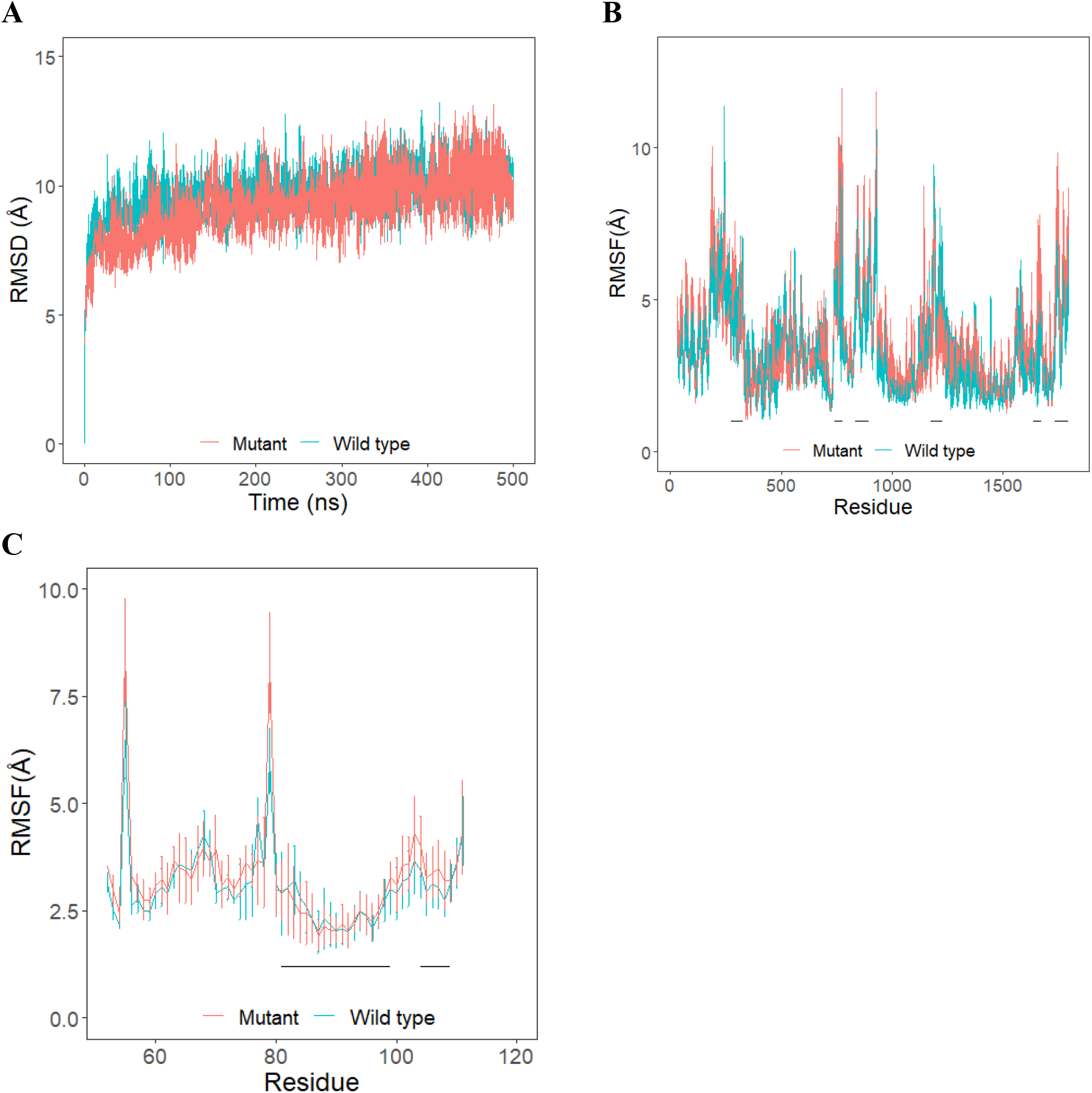
Root mean square deviation (RMSD) and root mean square fluctuation (RMSF) of protein Cα atoms with respect to the initial structure obtained from three independent runs. Results from three simulation runs of each system are plotted as mean ± SD. A) RMSD of wild-type and mutant IGF-1 – IGF-1R complexes; B) RMSF of Cα atoms of IGF-1R protein in the wild-type and mutant IGF-1 – IGF-1R complexes; C) RMSF of Cα atoms of IGF-1 protein in the wild-type and mutant IGF-1 – IGF-1R complexes. Loop and helical regions of each protomer of IGF-1R and IGF-1 binding regions that make contact with IGF-1R are identified with black bars.

### Comparison of regional fluctuations in the wild-type and mutant IGF-1 – IGF-1R complexes

To observe and compare backbone stability and fluctuations of the two complexes, root mean square fluctuation (RMSF) of backbone Cα atoms were measured and plotted **(Figure 3B and Supplementary Figures S3A and S3B)**. IGF-1’s primary binding site is composed of L1 residues Glu294, Glu333 and Ly336, and α-CT residues 727-741 of IGF-1R. The CR domain residues 276-328, residues 739-779 and 836-895 comprised of loop and helical regions of each protomer fluctuated more compared to the rest of the protein structure, while residues 329-734 of both protomers exhibited limited fluctuations **(Figure 3B)**. However, the mutant exhibited more fluctuation in these regions when compared to wild-type **(Figure 3B)**. Importantly, regions around the binding site residues also fluctuated more in the mutant **(Supplementary Figure S4)**.

The fluctuations of IGF-1 when bound to IGF-1R dimer structures were evaluated by assessing the RMSF of backbone Cα atoms of IGF-1. Overall, the IGF-1 backbone exhibited higher fluctuations in the mutant complex system, when compared to wild-type systems **(Figure 3C)**. Of all the IGF-1 residues, the loop regions spanning residues 54-57 and 76-79 were identified as very flexible, displaying elevated fluctuations across all systems **(Figure 3C)**. Interestingly, residues at the interfacial region of IGF-1 (81-99 and 104-109) showed less fluctuation in wild-type as compared to mutant **(Figure 3C)**. Additionally, IGF-1 residues Phe73, Pro87, Ile91, Val92, Asp93, Glu94 and Cys96, which are known to interact with IGF-1R, exhibited reduced fluctuation in wild-type **(Supplementary Figure S5)**. The more fluctuations observed in mutant IGF-1 suggested that *IGF-1*:p.Ile91Leu likely altered the binding of IGF-1 to IGF-1R compared to the wild-type.

### Interfacial residue contact duration differs substantially between wild-type IGF-1 - IGF-1R and mutant IGF-1 - IGF-1R complexes

During the simulations, several intermolecular contacts such as hydrophobic interactions, hydrogen bonds, salt bridges, π–π and cation–π interactions were observed to form, break, and reform. There were some interactions that lasted longer than others. The residues of wild-type and mutant IGF-1 that exhibited consistent interactions with IGF-1R are shown in **Figure 4**. The contact duration of intermolecular interactions between wild-type or mutant IGF-1 and IGF-1R interfaces, as well as the dynamics of each interaction throughout the duration of the simulation trajectories, are demonstrated in **Supplementary Table S4**.

**Figure 4.**
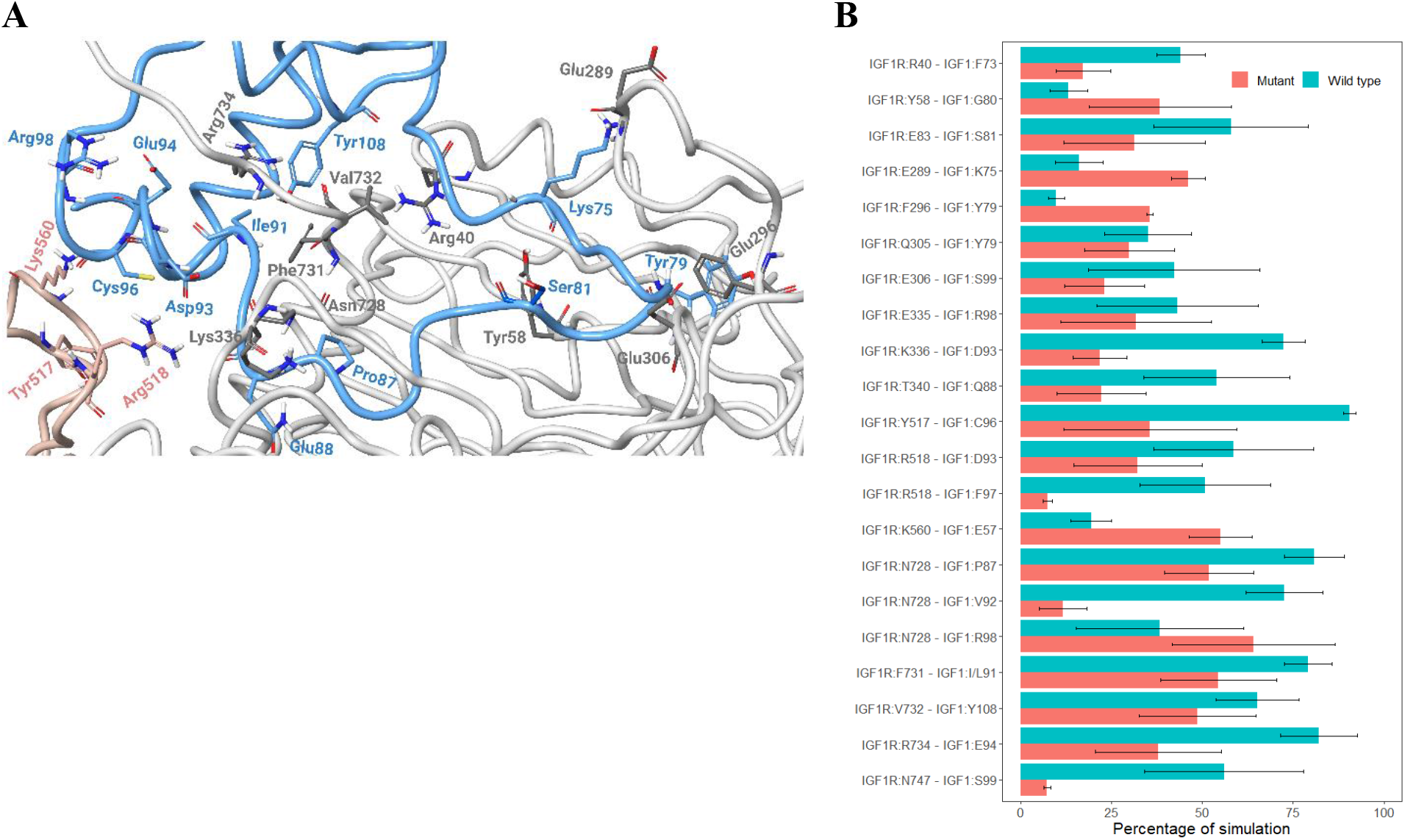
A) Enlarged view of the binding interface of IGF-1R (grey and pink) bound to IGF-1 (blue); B) The percentage of simulation time during which intermolecular contacts were retained between IGF-1R and IGF-1 interacting residues. Results from three simulation runs of each system are plotted as mean ± SEM.

In the wild-type simulation runs, IGF-1 residues Pro87 and Val92 formed consistent interactions with α-CT Asn728, while Lys75 and Tyr79 formed weak intermittent interactions with Glu289 and Phe296 residues of IGF-1R. Other essential α-CT residues, Val732 and Arg734, also formed more sustained interaction with wild-type IGF-1 Tyr108 and Glu94, respectively, compared to mutant IGF-1 **(Figure 4B)**. Additionally, IGF-1 residues Pro87 and Gln88 have been shown previously to form interactions with Phe725 and Ser729^18^. We, however, observed a stable interaction between Pro87 and Asn728 in wild-type only. Moreover, our findings differed from previous reports in that we did not observe Gln88 binding with Phe725 and Ser729 of the IGF-1R. Instead, Gln88 on IGF-1 consistently interacted with Thr340 in the wild-type, but not in the mutant.

In the wild-type IGF-1, Phe73 side chains underwent a rotameric rearrangement and were found buried in a hydrophobic pockets formed by Arg40, Leu63 and Phe731. Such a rearrangement helped Phe73 to form consistent interactions with Arg40. Interestingly, this interaction was observed to be weaker in the mutant compared to the wild-type **(Figure 4B)**. Notably, Phe71, Tyr72 and Phe73 have been shown to be important for IGF-1R binding^32^ and the rotameric rearrangement of Phe71 and Phe73 upon binding of IGF-1 to IGF-1R has previously been reported^18^. Furthermore, Asp93 of IGF-1 has been shown to exist near α-CT Asn724 as indicated in a cryo-EM structural study^11^. However, we have not observed this interaction in any of our simulation runs. Instead, Asp93 formed the more stable salt bridge and hydrogen bond with Lys336 and Arg518 in the wild-type compared to the mutant. Although, IGF-1 Val92 has been reported to make contacts with α-CT Asn724 and Asn728, our studies showed that Val92 formed a stable hydrogen bond with Asn728 in the wild-type IGF-1 as compared to mutant **(Figure 4)**. A mutagenesis study has illustrated the importance of Val92 in binding of IGF-1 to IGF-1R^33^. Additionally, Glu94 exhibited a stable interaction with IGF-1R residue Arg734 in the wild-type IGF-1 compared to mutant. A mutagenesis study has shown the essential role of Arg734 in the binding of IGF-1 with IGF-1R^34^.

IGF-1 C-terminal residues are known to be critical in maintaining optimal binding to IGF-1R. A replacement of C-terminal region with additional glycine residues have resulted in 30 -fold decrease in IGF-1 affinity for IGF-1R^35^. Here, IGF-1 C-terminal residues Ser99 and Tyr108 formed more consistent interactions with Asn747 and Val732 of IGF-1R in the wild-type compared to the mutant **(Figure 4B)**.

A small secondary binding site for IGF-1 within the active IGF-1R dimer has recently been reported^11^. It is mainly composed of loop regions of the FnIII-1′ domain. Residues 513-518 and Lys560 of FnIII-1′ form this secondary binding subsite. The IGF-1R residue Tyr517 formed interaction with Cys96 and Arg518 with Asp93 and Phe97. These interactions were noted to be more stable in wild-type runs **(Figure 4B and Supplementary Table S4)**. Interestingly, Lys560 formed a much more stable salt bridge with IGF-1 residue Glu57 in the mutant runs compared to wild-type **(Figure 4B)**. A previous mutagenesis study reported that residues in a subsite (513-518), particularly Tyr517 and Arg518, were important for IGF-1 optimal binding, whereas residues around Lys560 had no effect on IGF-1 dependent IGF-1R activation^11^. However, the precise role of Lys560 has not been established yet.

Interestingly, mutant variant *IGF-1*:p.Ile91Leu is located at the binding interface of IGF-1. Structural studies have indicated that Ile91 residue makes contact with His727, Asn728 and Phe731. The wild-type IGF-1 protein formed relatively more sustained interactions with Phe731, compared to mutant IGF-1 **(Figure 4B)**. It is perceivable that this interaction helped nearby residues of IGF-1, particularly Val92, Asp93 and Glu94 to form stable interactions with IGF-1R interfacial residues **(Figure 4)**.

### Wild-type IGF-1 bound more stably to IGF-1R

An analysis of intermolecular interactions demonstrated that IGF-1R complexes formed a greater number of interactions with the wild-type IGF-1 compared to the mutant IGF-1 **(Figure 4 and Supplementary Table S4)**. This finding suggested that compared to the mutant IGF-1, the wild-type IGF-1 may have greater affinity for the IGF-1R. To examine the energetic contributions, the free energy of binding (ΔG_bind_) was compared between mutant and wild-type IGF-1 bound to IGF-1R, calculated using the molecular mechanics-generalized Born surface area (MM-GBSA) approach based on frames extracted every 2.5 ns from all MD simulations. All wild-type simulations demonstrated substantially higher ΔG_bind_ values than the mutant runs **(Table 1)**. A higher free energy of binding indicated a stronger binding affinity. Hydrophobic contributions to ΔG_bind_ were also lower for mutant simulations compared to wild-type. The complete count of intermolecular hydrogen bonds between the two complexes was also tracked during the simulations. The wild-type complexes exhibited a greater number of hydrogen bonds (mean ± SD for three simulations: 23.88 ± 3.50, 25.85 ± 3.65, 21.88 ± 4.02) compared to the mutant (20.65 ± 4.01, 24.06 ± 4.46, 18.64 ± 3.28) in all simulation runs **(Supplementary Figure S6)**. This would also be expected to enhance the binding affinity between wild-type IGF-1 and IGF-1R.

**Table 1.** MM-GBSA based free energy of binding of IGF-1 at site 1 and site 2 of IGF-1R.

| Run | Site 1 |  | Site 2 |  |
| --- | --- | --- | --- | --- |
| | Wild-type (Mean ± SD) | Mutant (Mean ± SD) | Wild-type (Mean ± SD) | Mutant (Mean ± SD) |
| 1 | -266.39 ± 35.52 | -212.66 ± 46.48 | -12.85±6.16 | -18.55±21.14 |
| 2 | -258.52 ± 26.72 | -234.54 ± 31.94 | -5.90± 0.84 | -7.3±1.41 |
| 3 | -231.77 ± 20.47 | -223.76 ± 22.81 | -7.65±0.49 | -7.41±1.27 |

It has long been known that two identical binding sites exist on the IGF-1R dimer in apo state, indicating that IGF-1 can bind with similar probability to either one of the two sites ^11,17^. Upon IGF-1 binding to either site on the IGF-1R dimer, IGF-1R undergoes structural rearrangement and subsequently the L1 domain, α-CT (ligand bound), and the bound IGF-1 ascend towards the upper section of the IGF-1R dimer and is obligatory for the binding of a second IGF-1 molecule^11^. This is the known phenomenon of negative cooperativity. Based on this, we hypothesized that mutant IGF-1R bound to one IGF-1 may alter the binding of a second IGF-1 molecule. To test this, the last frame of each simulation run was extracted and a second IGF-1 molecule was docked into the second binding site of the IGF-1R. Both the wild-type and the mutant IGF-1 bound complexes exhibited very weak binding of the second IGF-1 to the IGF-1R, as evidenced by their respective binding affinity scores **(Table 1)**. This suggested that mutant IGF-1 did not alter the negative cooperativity of IGF-1R.

### Mutant alters IGF-1R inter-protomer interactions

We also investigated the impact of mutant IGF-1 on dynamics of IGF-1R inter-protomer interactions. The inter-protomer interactions are shown in **Figure 5 and Supplementary Table S5**. Specific residues within L1–FnIII-2′ have been demonstrated to be crucial for IGF-1R dimerization^18^. In both wild-type and mutant simulations, Glu177 and Glu188 residues of L1 formed similar interactions with Arg671 and Lys665 of FnIII-2′, respectively **(Figure 5A and Supplementary Figure S7)**. A network of inter-protomer interactions showed similar binding stability throughout the simulation runs of IGF-1R bound with the wild-type and the mutant IGF-

**Figure 5.**
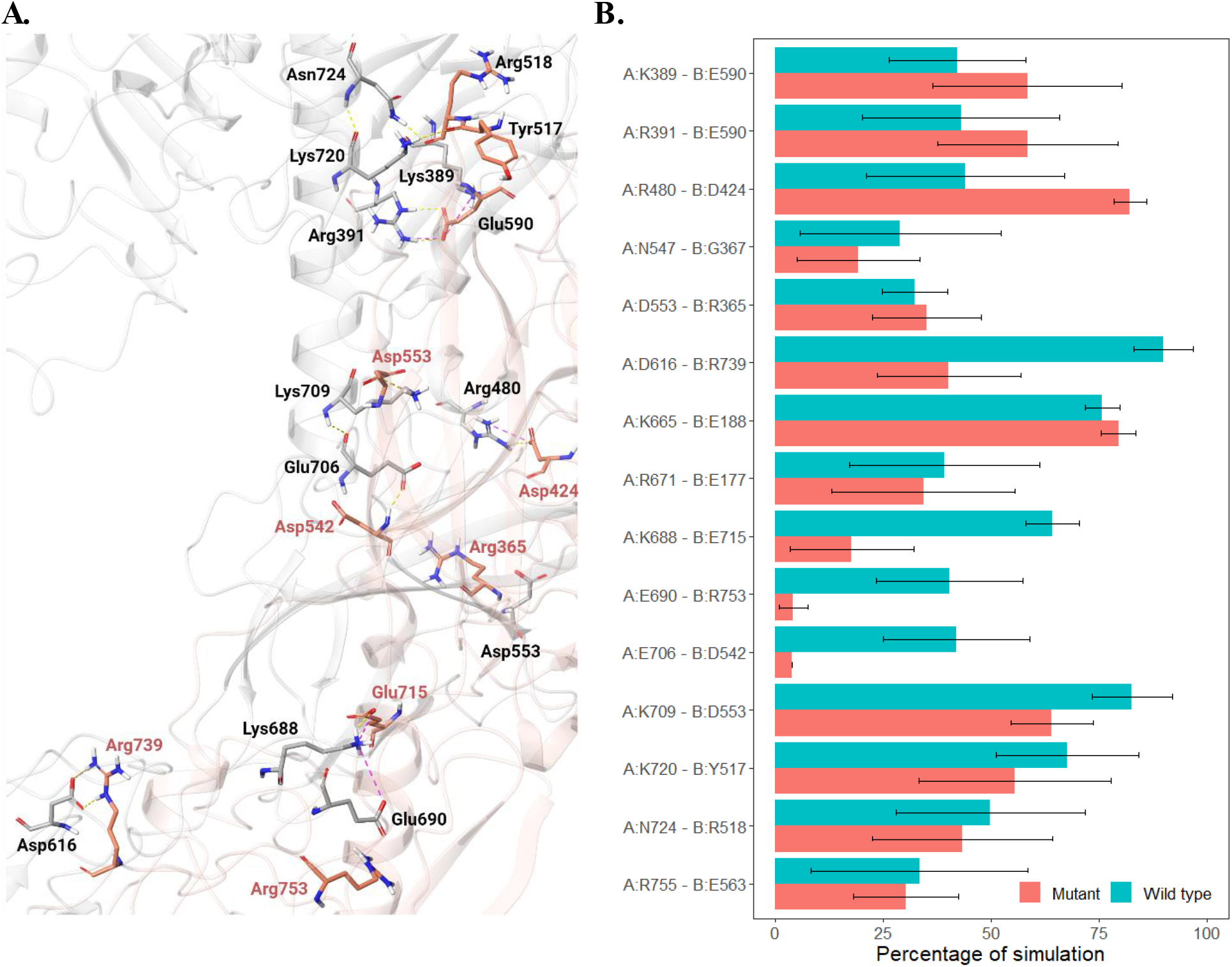
Binding interface of chain A and Chain B of IGF-1R. A) Enlarged binding pose showing the residues that interact in the interface; B) The percentage of simulation time during which intermolecular contacts were retained between chain A and chain B interacting residues of IGF-1R dimer. Results from three simulation runs of each system are plotted as mean ± SEM. Hydrogen bonds and salt bridges are represented by yellow and pink dotted lines, respectively.

1. For instance, in all simulation runs, chain A residues Asp553, Lys720, Asn724, and Arg755 showed similar interactions stability with chain B residues Arg365, Tyr517, Arg518, and Glu563, respectively (**Figure 5 and Supplementary Table S5)**. However, A:Lys389-B:Glu590, A:Arg391-B:Glu590, A:Arg480-B:Asp424, A:Asp616-B:Arg739, A:Lys688-B:Glu715, A:Glu690-B:Arg753, A:Glu706-B:Asp542 and A:Lys709 – B:Asp553 inter-protomer interactions varied substantially between wild-type and mutant runs **(Figure 5)**. Overall, compared to the mutant, the wild-type simulations resulted in the formation of slightly more stable interactions between two chains of the IGF-1R. This indicated that mutant IGF-1 caused a subtle conformational change in the IGF-1R. This subtle rearrangement of IGF-1R protomers likely changed the configuration of IGF-1 binding pocket and weakened the binding of the mutant IGF-1 to IGF-1R.

## Discussion

This study provided insights into two IGF-1 gene coding variants discovered in individuals with exceptional longevity. We described the structural and functional impact of IGF-1:p.Ile91Leu by characterizing the stability of interactions that define the IGF-1R – IGF-1 interface. Utilization of extended MD simulations demonstrated that compared to the wild-type IGF-1, the IGF-1:p.Ile91Leu variant resulted in weaker interactions between IGF-1 and its receptor, likely attenuating IGF-1R activation. Additionally, we identified the *IGF-1*:p.Ala118Thr variant, which was significantly associated with lower levels of IGF-1 in our longevity cohort. The latter variant may result in lower circulating IGF-1 level due to its location near an *IGF-1* gene E-peptide region, which is typically removed during the post-translational processing of the IGF-1 precursor protein. Overall, our results suggest that compared to wild-type IGF-1, the activation of IGF-1R and subsequent downstream signaling mediated by mutant IGF-1s would be expected to produce attenuated effects. Given the previously identified role of reduced insulin/IGF-1 signaling in models of longevity, our findings provide additional evidence for the potential role of these gene variants and reduced IGF-1 signaling in human longevity.

Previous longevity-focused GWAS studies have not identified signals in *IGF-1* gene, despite the conserved roles of insulin/IGF-1 system in longevity. This is not entirely surprising since investigations of common genetic variants that occur at frequencies of >5% in the general population, to study the uncommon event of exceptional longevity that generally occurs at a rate of <1% in the population, are likely to miss the rare longevity-associated genotypes^36^. In this study, we attempted to find all coding variants in the *IGF-1* gene in our longevity cohort. Interestingly, we found only two rare coding variants (MAF ≤0.01), supporting the idea that IGF-1 is highly conserved across species. Moreover, rare variants association studies to date have generally been underpowered to detect their effects on phenotype^37^. One approach to overcome this challenge is to perform GWAS in very large longevity cohorts, which are currently unavailable. Alternatively, one can focus on rare variants that may be more represented among individuals with longevity. Studies have demonstrated the functional impacts of longevity specific rare coding variants at the single gene level, in genes such as IGF-1R^5^, SIRT6^38^, APOC3^39^, as well as others. For instance, a previous study identified the *IGF-1R*.p.Ala67Thr variant in only two centenarians^5^. This variant was shown to cause decreased IGF-1R activation, possibly by weakening its binding to IGF-1. Similarly, a recent study found two rare coding variants (rs183444295 and rs201141490) in *SIRT6* among centenarians^38^. Interestingly, the *SIRT6* variants were found to strongly suppress LINE1 retrotransposons, boost DNA double-strand break repair, and more effectively eradicate cancer cells compared to the wild-type. These studies indicate the importance of identifying and establishing the molecular mechanisms of longevity associated rare coding variants. Understanding the mechanisms of rare longevity-associated variants found in individuals with exceptional longevity is of paramount importance in advancing our knowledge of how genes that carry these variants regulate downstream signaling of pro-longevity pathways and could serve as promising gerotherapeutic drug targets.

The *IGF-1* gene significantly impacts growth and development. A genetic variant located in the promotor region of the *IGF-1* gene has been shown to be associated with small size in dogs^40^. In mice, a synonymous mutation in *IGF-1* significantly affected both the expression and biological functions of IGF-1^41^. Short stature and reduced binding affinity of IGF-1 to IGF-1R have also been reported in families with coding variants in the *IGF-1* gene^6,42^. However, to the best of our knowledge, *IGF-1* coding variants have not been previously identified in humans with longevity. We identified two likely functional coding variants, Ile91Leu and Ala118Thr, in *IGF-1* in a heterozygous state that were not associated with adult maximal height, likely because they result in partial reduction of IGF-1 function. Similarly, centenarian specific variants that induced only partial loss of function have been identified in IGF-1R^5^. Interestingly, we found the *IGF-1*:p.Ile91Leu variant in two centenarians, located at the binding interface of IGF-1R – IGF-1. Although, from a physiochemical perspective the substitution of isoleucine to leucine is not expected to be functionally significant, this change has previously been shown to alter protein-protein interactions and enzyme activity in other genes^43-45^. Potential functional effects of missense substitutions are illustrated by a prior study, in which the assessment of various IGF-1 analogs revealed that [His95]-IGF-1 and [Gln95]-IGF-1 exhibited significantly reduced binding affinities for IGF-1R that resulted in diminished activation of IGF-1R compared to wild-type^46^. A similar pattern was also noted in another study wherein IGF-1 analogs that exhibited weaker binding affinity demonstrated reduced activation of IGF-1R^10^. This implies that a higher binding affinity of IGF-1 does lead to a more robust activation of IGF-1R. Moreover, a specific conformational change at the cytoplasmic end of IGF-1R upon IGF-1 binding is crucial to generate optimal downstream signaling. In this study, *IGF-1*:p.Ile91Leu demonstrated reduced binding affinity with IGF-1R at the extracellular binding site compared to wild-type. Consistent with earlier studies, it is likely that this variant will induce a change in the conformation of IGF-1R at the cytoplasmic end, potentially reducing its activation. Diminished IGF-1R signaling has consistently been shown to extend lifespan in multiple model organisms^18,47,48^, including humans^49^, where individuals with exceptional longevity and IGF-1R coding variants exhibited reduced activity of IGF-1R and IGF-1 induced AKT phosphorylation^5^.

The epidemiological studies that focused on assessing the association of circulating IGF-1 levels with life-span and health-span have shown mixed results^49^. Studies in longevity cohorts have reported positive associations between higher IGF-1 with all-cause mortality and age-related diseases^50-52^. Conversely, other studies, mostly involving younger populations, have indicated the reverse: elevated IGF-1 levels were linked to a decreased risk of disease and mortality^53,54^. However, a recent large-scale study involving nearly 450,000 UK biobank participants showed that older adults with higher IGF-1 levels had greater risk of mortality and age-related diseases, indicating that lower IGF-1 levels were beneficial for their survival^55^. In our longevity cohort, carriers of *IGF-1*:p.Ala118Thr had significantly lower levels of IGF-1, compared to non-carriers (Figure 1B). Interestingly, a synonymous variant in exon 4 of the IGF-1 gene has previously been shown to reduce the expression, secretion, stability, and half-life of IGF-1 in mice. In our study, *IGF-1*:p.Ala118Thr was located at the intersection of Exon 4 and the N-terminal sequence of the E-peptides (pro-peptides) of IGF-1 (Figure 1A). Moreover, this variant falls within a unique pentabasic motif (Lys113-Arg125) where post-translational cleavage of pro-IGF-1 polypeptides generally occurs. This cleavage has been shown to regulate the expression, stability, release and bioavailability of IGF-1^20,56,57^. Thus, this variant may potentially modify the binding motif involved in the cleavage of the carboxyl-terminal E domain from the pro-IGF-, resulting in lower IGF-1 level. Lower IGF-1 levels may in turn lead to reduced IGF-1R signaling^6,55^, which may be beneficial for longevity. This variant may have an allosteric effect on IGF-1binding, potentially reducing its interaction with the IGF-1R. However, the *IGF-1*:p.Ala118Thr variant dependent effect on the expression, stability, release, and bioavailability of IGF-1 molecule is also possible.

In summary, this study identified two rare functional coding variants *IGF-1*:p.Ile91Leu and *IGF-1*:p.Ala118Thr which likely impact the IGF-1 induced downstream signaling of IGF-1R. Our findings suggest that *IGF-1*:p.Ile91Leu and *IGF-1*:p.Ala118Thr variants attenuate IGF-1R activity, potentially via reduced binding of IGF-1 to IGF-1R and by diminishing the circulatory levels of IGF-1, respectively. These results provide evidence that the rare *IGF-1* variants identified in cohorts with exceptional longevity may contribute to extended lifespan via attenuation of IGF-1 signaling.

## Methods

### Recruitment of study participants

The participants in this study were Ashkenazi Jews from two well-characterized longevity cohorts, the Longevity Genes Project (LGP) and the LonGenity study, which have been recruited and characterized at the Albert Einstein College of Medicine. The longevity cohorts consisted of individuals with exceptional longevity (centenarians) age ≥95 years, offspring of individuals with exceptional longevity (offspring), defined as having at least one parent who lived to 95 years or older, and individuals without parental history of exceptional longevity (controls), defined as not having a parent that survived beyond 95 years of age. Both LGP and LonGenity studies were approved by the institutional review board (IRB) of Albert Einstein College of Medicine^58-60^ (approval numbers 1998-125 and 2007-272, respectively) and were performed in compliance with the Declaration of Helsinki. Written informed consent was obtained from all subjects. All experimental protocols were approved by IRB of Albert Einstein College of Medicine (approval numbers 1998-125 and 2007-272, respectively).

### Whole exome sequencing and functional variant identification

Whole exome sequencing (WES) of 2,521 subjects was carried out at the Regeneron Genetics Center (RGC). The pipeline adopted for sample preparation and WES has been previously described^61^. GRCh38 human genome assembly was used for variant calling. Individuals with low sequencing coverage (less than 80% of bases with coverage ≥ 20x), call rate < 0.9, and discordant sex were excluded. SNPs were removed if they had the read depth (DP) < 7 (DP < 10 for insertions/deletions (INDEL)), alternative Allele Balance less than a cutoff (≤ 15% for SNP, ≤ 20% for INDEL), and Hardy–Weinberg equilibrium deviated from an *χ*2-test P < 1 × 10^−6^. Variants with missing rates < 0.01 in the study cohort were used for further analysis. In this study, we focused on rare variants with minor allele frequencies <1% in our cohort. The functional nature of the variants was predicted using combined annotation dependent depletion (CADD) score^27^. It is a widely used method to predict the variant’s deleteriousness. Variants with CADD score ≥ 20 were considered functional. Overall, 2,487 subjects and 2 variants in the *IGF-1* gene passed all the thresholds.

### Protein modeling and molecular dynamics (MD) simulations

The three dimensional (3D) dimeric crystal structure of human IGF-1R was retrieved from the Protein Data Bank (PDB ID: 6JK8). The protein structure was visualized and prepared for docking by using Schrödinger Maestro 2023-2 (Schrödinger, LLC, NY). The structure was first pre-processed using the Protein Preparation Wizard (Schrödinger, LLC, NY). The protein preparation stage included proper assignment of bond order, adjustment of ionization states, orientation of disorientated groups, creation of disulphide bonds, removal of unwanted water molecules, metal and co-factors, capping of the termini, assignment of partial charges, and addition of missing atoms and side chains using default protein preparation wizard tasks. Loops refinement and further structural verification was carried out using the protein refinement module of Schrödinger Prime using default settings. The missing hydrogen atoms were added, and standard protonation state at pH7 was used. The human wild-type and the mutant IGF-1 was docked in the binding site 1 of the IGF-1R using protein-protein docking suite (BioLuminate, Schrödinger, LLC, NY). IGF-1 protein was used as ligand and was docked starting from multiple random conformations. Ten representative docked protein-protein complexes were chosen following the clustering of the generated conformers. The binding poses of IGF-1 with IGF-1R were compared with the already reported structures. The best binding pose of wild-type and mutant IGF-1 with IGF-1R based on free energy of binding, was subjected to MD simulations. Mutant *IGF-1*:p.Ile91Leu was generated by employing computational point mutations using the residue mutation panel of Schrodinger Maestro. Structures of wild-type and mutant IGF-1 bound to IGF-1R were placed in large orthorhombic boxes of size 160 Å × 160 Å × 230 Å and solvated with single point charge (SPC) water molecules using the Desmond System Builder (Schrödinger, LLC, NY). An appropriate number of counterions were added to neutralize the simulation systems and salt concentration of 0.15 M NaCl was maintained. All-atom MD simulations were carried out using Desmond^62^. All calculations were performed using the OPLS forcefield. Prior to the start of the production run, all prepared simulation systems were subjected to Desmond’s default eight stage relaxation protocol. Both the wild-type and mutant IGF-1 bound to IGF-1R were simulated for 500 ns in triplicates using different sets of initial seed velocities. To maintain the pressure at 1 atm and temperature at 300 K during the simulation runs, the isotropic Martyna–Tobias–Klein barostat^63^ and the Nose– Hoover thermostat^64^ were used, respectively. A 9.0 Å cutoff was set for short-range interactions and the smooth particle mesh Ewald method (PME)^65^ was used to measure the long-range coulombic interactions. A time-reversible reference system propagator algorithm (RESPA) integrator was used with an inner time step of 2.0 fs and an outer time step 6.0 fs. Molecular Mechanics-Generalized Born Surface Area (MM-GBSA) method was employed to determine the free energy of binding of wild-type and mutant IGF-1 protein to IGF-1R using frames obtained from MD simulation trajectories. Frames were retrieved every 2.5 ns from each of the simulation runs and MM-GBSA based binding free energy was computed using Schrödinger Prime employing the VSGB 2.0 solvation model^66^. Protein-Protein docking was carried out to dock the additional IGF-1 molecule at the second binding site of IGF-1R (BioLuminate, Schrödinger, LLC, NY). For this, the last frame from each simulation run was extracted, and an additional IGF-1 molecule was docked into the second binding site of the IGF-1R. Three independent runs of protein-protein docking were performed on each structure. The top binding poses in each run were subjected to MM-GBSA to evaluate the binding free energy in an implicit solvent model. Simulation data was analyzed using packaged and in-house scripts. Graphs were plotted using R version 3.6.3 (https://www.r-project.org) and images of structures were generated using Schrödinger Maestro 2023-2 (Schrödinger, LLC, NY).

## Acknowledgements

This work was supported by R01AG061155 to SM and the American Federation for Aging Research/Glenn Foundation for Medical Research Postdoctoral Fellow grant to AA.

## Contributions

AA and SM conceived the idea. SM, NR, TG and SA enrolled the study participants and performed clinical examinations. AA performed the experiments. AA, SM, ZZ, EG, and NB performed the analysis. AA and SM wrote the manuscript.

## Data availability statement

The datasets generated during and/or analysed during the current study are available from the corresponding authors on reasonable request.

## References

1. Holzenberger, M. et al. IGF-1 receptor regulates lifespan and resistance to oxidative stress in mice. Nature 421, 182–187 (2003).

2. Bartke, A. Minireview: role of the growth hormone/insulin-like growth factor system in mammalian aging. Endocrinology 146, 3718–23 (2005).

3. Coschigano, K.T. et al. Deletion, but not antagonism, of the mouse growth hormone receptor results in severely decreased body weights, insulin, and insul in-like growth factor I levels and increased life span. Endocrinology 144, 3799–810 (2003).

4. Russell, S.J. & Kahn, C.R. Endocrine regulation of ageing. Nat Rev Mol Cell Biol 8, 681–91 (2007).

5. Suh, Y. et al. Functionally significant insulin-like growth factor I receptor mutations in centenarians. Proceedings of the National Academy of Sciences of the United States of America 105, 3438–3442 (2008).

6. Giacomozzi, C. et al. Novel Insulin-Like Growth Factor 1 Gene Mutation: Broadening of the Phenotype and Implications for Insulin Resistance. Journal of Clinical Endocrinology & Metabolism 108, 1355–1369 (2023).

7. Netchine, I. et al. Partial primary deficiency of insulin-like growth factor (IGF)-I activity associated with IGF1 mutation demonstrates its critical role in growth and brain development. J Clin Endocrinol Metab 94, 3913–21 (2009).

8. Bonapace, G., Concolino, D., Formicola, S. & Strisciuglio, P. A novel mutation in a patient with insulin-like growth factor 1 (IGF1) deficiency. J Med Genet 40, 913–7 (2003).

9. Walenkamp, M.J. et al. Homozygous and heterozygous expression of a novel insulin-like growth factor-I mutation. J Clin Endocrinol Metab 90, 2855–64 (2005).

10. Machácková, K. et al. Insulin-like Growth Factor 1 Analogs Clicked in the C Domain: Chemical Synthesis and Biological Activities. Journal of Medicinal Chemistry 60, 10105–10117 (2017).

11. Li, J., Choi, E., Yu, H.T. & Bai, X.C. Structural basis of the activation of type 1 insulin-like growth factor receptor. Nature Communications 10(2019).

12. Siddle, K. Molecular basis of signaling specificity of insulin and IGF receptors: neglected corners and recent advances. Front Endocrinol (Lausanne) 3, 34 (2012).

13. Smith, B.J. et al. Structural resolution of a tandem hormone-binding element in the insulin receptor and its implications for design of peptide agonists. Proc Natl Acad Sci U S A 107, 6771–6 (2010).

14. Whittaker, J. et al. Alanine scanning mutagenesis of a type 1 insulin-like growth factor receptor ligand binding site. J Biol Chem 276, 43980–6 (2001).

15. Whittaker, L., Hao, C., Fu, W. & Whittaker, J. High-affinity insulin binding: insulin interacts with two receptor ligand binding sites. Biochemistry 47, 12900–9 (2008).

16. Mynarcik, D.C., Yu, G.Q. & Whittaker, J. Alanine-scanning mutagenesis of a C-terminal ligand binding domain of the insulin receptor alpha subunit. J Biol Chem 271, 2439–42 (1996).

17. Kavran, J.M. et al. How IGF-1 Activates its Receptor. Elife 3(2014).

18. Xu, Y.B. et al. How ligand binds to the type 1 insulin-like growth factor receptor. Nature Communications 9(2018).

19. Zhang, X. et al. Visualization of Ligand-Bound Ectodomain Assembly in the Full-Length Human IGF-1 Receptor by Cryo-EM Single-Particle Analysis. Structure 28, 555–561 e4 (2020).

20. Philippou, A., Maridaki, M., Pneumaticos, S. & Koutsilieris, M. The complexity of the IGF1 gene splicing, posttranslational modification and bioactivity. Mol Med 20, 202–14 (2014).

21. Wang, S.Y. et al. A synonymous mutation in IGF-1 impacts the transcription and translation process of gene expression. Mol Ther Nucleic Acids 26, 1446–1465 (2021).

22. Song, X.T. et al. Molecular cloning, expression, and functional features of IGF1 splice variants in sheep. Endocr Connect 10, 980–994 (2021).

23. Freitas, E.D.S. et al. Lower muscle protein synthesis in humans with obesity concurrent with lower expression of muscle IGF1 splice variants. Obesity (Silver Spring) 31, 2689–2698 (2023).

24. Chrudinova, M. et al. A viral insulin-like peptide inhibits IGF-1 receptor phosphorylation and regulates IGF1R gene expression. Mol Metab 80, 101863 (2024).

25. De Vivo, M., Masetti, M., Bottegoni, G. & Cavalli, A. Role of Molecular Dynamics and Related Methods in Drug Discovery. Journal of Medicinal Chemistry 59, 4035–4061 (2016).

26. Wingler, L.M., McMahon, C., Staus, D.P., Lefkowitz, R.J. & Kruse, A.C. Distinctive Activation Mechanism for Angiotensin Receptor Revealed by a Synthetic Nanobody. Cell 176, 479-+ (2019).

27. Kircher, M. et al. A general framework for estimating the relative pathogenicity of human genetic variants. Nat Genet 46, 310–5 (2014).

28. Rotwein, P. Diversification of the insulin-like growth factor 1 gene in mammals. Plos One 12(2017).

29. Thompson, J.D., Higgins, D.G. & Gibson, T.J. Clustal -W -Improving the Sensitivity of Progressive Multiple Sequence Alignment through Sequence Weighting, Position-Specific Gap Penalties and Weight Matrix Choice. Nucleic Acids Research 22, 4673–4680 (1994).

30. Savva, L. & Platts, J.A. Computational investigation of copper-mediated conformational changes in alpha-synuclein dimer. Phys Chem Chem Phys 26, 2926–2935 (2024).

31. Shinwari, K. et al. In-silico assessment of high-risk non-synonymous SNPs in ADAMTS3 gene associated with Hennekam syndrome and their impact on protein stability and function. Bmc Bioinformatics 24(2023).

32. Cascieri, M.A. et al. Mutants of Human Insulin-Like Growth Factor-I with Reduced Affinity for the Type-1 Insulin-Like Growth-Factor Receptor. Biochemistry 27, 3229–3233 (1988).

33. Denley, A. et al. Structural and functional characteristics of the Val44Met insulin-like growth factor I missense mutation: correlation with effects on growth and development. Mol Endocrinol 19, 711–21 (2005).

34. Kertisová, A. et al. Insulin receptor Arg717 and IGF-1 receptor Arg704 play a key role in ligand binding and in receptor activation. Open Biology 13(2023).

35. Bayne, M.L. et al. The C-Region of Human Insulin-Like Growth-Factor (Igf)-I Is Required for High-Affinity Binding to the Type-1 Igf Receptor. Journal of Biological Chemistry 264, 11004–110 08 (1989).

36. Milman, S. & Barzilai, N. Discovering Biological Mechanisms of Exceptional Human Health Span and Life Span. Cold Spring Harb Perspect Med 13(2023).

37. Barton, A.R., Sherman, M.A., Mukamel, R.E. & Loh, P.R. Whole-exome imputation within UK Biobank powers rare coding variant association and fine-mapping analyses. Nature Genetics 53, 1260-+ (2021).

38. Simon, M. et al. A rare human centenarian variant of SIRT6 enhances genome stability and interaction with Lamin A. EMBO J 42, e113326 (2023).

39. Atzmon, G. et al. Lipoprotein genotype and conserved pathway for exceptional longevity in humans. PLoS Biol 4, e113 (2006).

40. Sutter, N.B. A single allele is a major determinant of small size in dogs (vol 316, pg 112, 2007). Science 316, 1284–1284 (2007).

41. Wang, S.Y. et al. A synonymous mutation inimpacts the transcription and translation process of gene expression. Molecular Therapy-Nucleic Acids 26, 1446–1465 (2021).

42. Walenkamp, M.J.E. et al. Homozygous and heterozygous expression of a novel insulin-like growth factor-I mutation. Journal of Clinical Endocrinology & Metabolism 90, 2855–2864 (2005).

43. Wu, E. et al. A conservative isoleucine to leucine mutation causes major rearrangements and cold-sensitivity in KlenTaq1 DNA polymerase. Faseb Journal 28(2014).

44. Sitbon, M. et al. Substitution of Leucine for Isoleucine in a Sequence Highly Conserved among Retroviral Envelope Surface Glycoproteins Attenuates the Lytic Effect of the Friend Murine Leukemia-Virus. Proceedings of the National Academy of Sciences of the United States of America 88, 5932–5936 (1991).

45. He, L. et al. Single methyl groups can act as toggle switches to specify transmembrane Protein-protein interactions. Elife 6(2017).

46. Machácková, K. et al. Converting Insulin-like Growth Factors 1 and 2 into High-Affinity Ligands for Insulin Receptor Isoform A by the Introduction of an Evolutionarily Divergent Mutation. Biochemistry 57, 2373–2382 (2018).

47. Kenyon, C., Chang, J., Gensch, E., Rudner, A. & Tabtiang, R. A C-Elegans Mutant That Lives Twice as Long as Wild-Type. Nature 366, 461–464 (1993).

48. Holzenberger, M. et al. IGF-1 receptor regulates lifespan and resistance to oxidative stress in mice. Nature 421, 182–7 (2003).

49. Milman, S., Huffman, D.M. & Barzilai, N. The Somatotropic Axis in Human Aging: Framework for the Current State of Knowledge and Future Research. Cell Metabolism 23, 980–989 (2016).

50. Milman, S. et al. Low insulin-like growth factor-1 level predicts survival in humans with exceptional longevity. Aging Cell 13, 769–771 (2014).

51. van der Spoel, E. et al. Association analysis of insulin-like growth factor-1 axis parameters with survival and functional status in nonagenarians of the Leiden Longevity Study. Aging (Albany NY) 7, 956–63 (2015).

52. Zhang, W.B. et al. Insulin-like Growth Factor-1 and IGF Binding Proteins Predict All-Cause Mortality and Morbidity in Older Adults. Cells 9(2020).

53. Bourron, O. et al. Impact of age-adjusted insulin-like growth factor 1 on major cardiovascular events after acute myocardial infarction: results from the fast-MI registry. J Clin Endocrinol Metab 100, 1879–86 (2015).

54. Friedrich, N. et al. Mortality and serum insulin-like growth factor (IGF)-I and IGF binding protein 3 concentrations. J Clin Endocrinol Metab 94, 1732–9 (2009).

55. Zhang, W.B., Ye, K., Barzilai, N. & Milman, S. The antagonistic pleiotropy of insulin-like growth factor 1. Aging Cell 20, e13443 (2021).

56. Hede, M.S. et al. E-peptides control bioavailability of IGF-1. PLoS One 7, e51152 (2012).

57. Annibalini, G. et al. The intrinsically disordered E-domains regulate the IGF-1 prohormones stability, subcellular localisation and secretion. Sci Rep 8, 8919 (2018).

58. Barzilai, N. et al. Unique lipoprotein phenotype and genotype associated with excepti onal longevity. Jama-Journal of the American Medical Association 290, 2030–2040 (2003).

59. Ismail, K. et al. Compression of Morbidity Is Observed Across Cohorts with Exceptional Longevity. Journal of the American Geriatrics Society 64, 1583–1591 (2016).

60. Gubbi, S. et al. Effect of Exceptional Parental Longevity and Lifestyle Factors on Prevalence of Cardiovascular Disease in Offspring. Am J Cardiol 120, 2170–2175 (2017).

61. Lin, J.R. et al. Rare genetic coding variants associated with human longevity and protection against age-related diseases. Nat Aging 1, 783–794 (2021).

62. Bowers, K.J. et al. Scalable algorithms for molecular dynamics simulations on commodity clusters. in Proceedings of the 2006 ACM/IEEE Conference on Supercomputing 84-es (2006).

63. Martyna, G.J., Tobias, D.J. & Klein, M.L. Constant pressure molecular dynamics algorithms. The Journal of chemical physics 101, 4177–4189 (1994).

64. Martyna, G.J., Klein, M.L. & Tuckerman, M. Nosé–Hoover chains: The canonical ensemble via continuous dynamics. The Journal of chemical physics 97, 2635–2643 (1992).

65. Essmann, U. et al. A smooth particle mesh Ewald method. The Journal of chemical physics 103, 8577–8593 (1995).

66. Li, J. et al. The VSGB 2.0 model: a next generation energy model for high resolution protein structure modeling. Proteins: Structure, Function, and Bioinformatics 79, 2794–2812 (2011).

