## Supplemental file 1 for "Identification of functional rare coding variants in IGF-1 gene in humans with exceptional longevity"

### Supplementary Materials

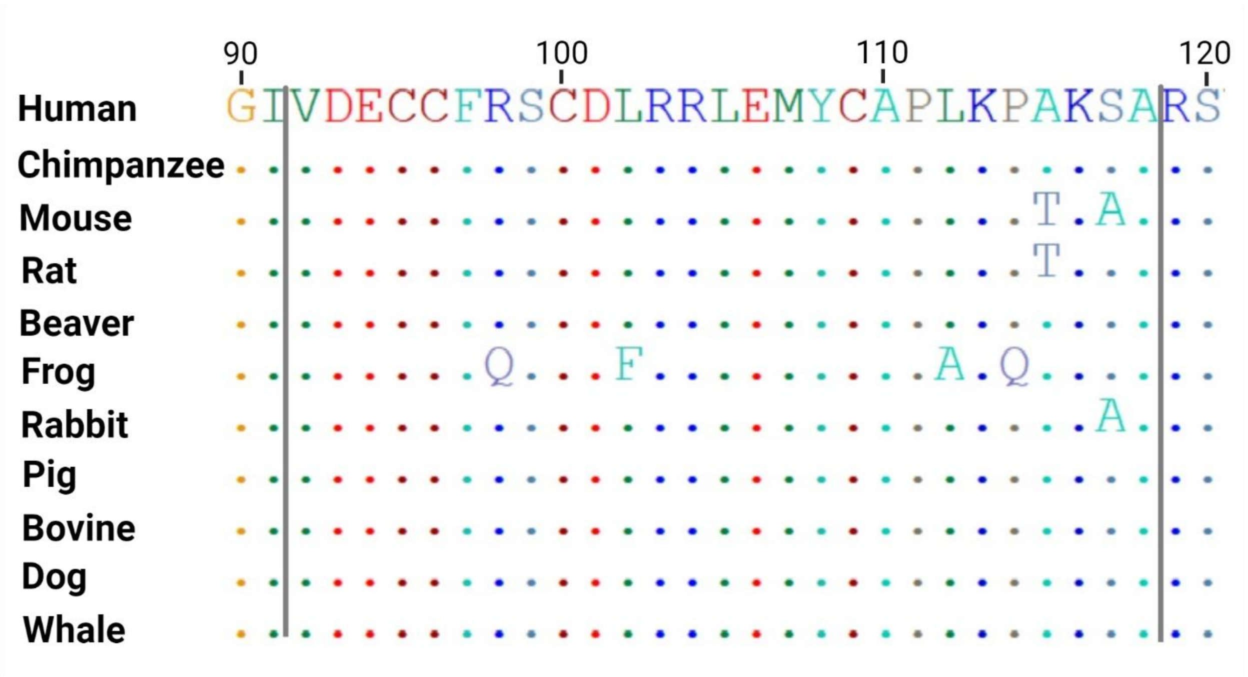


**Figure S1.** Multiple sequence alignment of thirty amino acids centered on the identified variants obtained from different mammals. Positions where sequences match the consensus are indicated with a standard dot, while mismatches are highlighted with alphabet. A) IGF-1, Ile91Leu and Ala118Thr.

**A.** **B.**

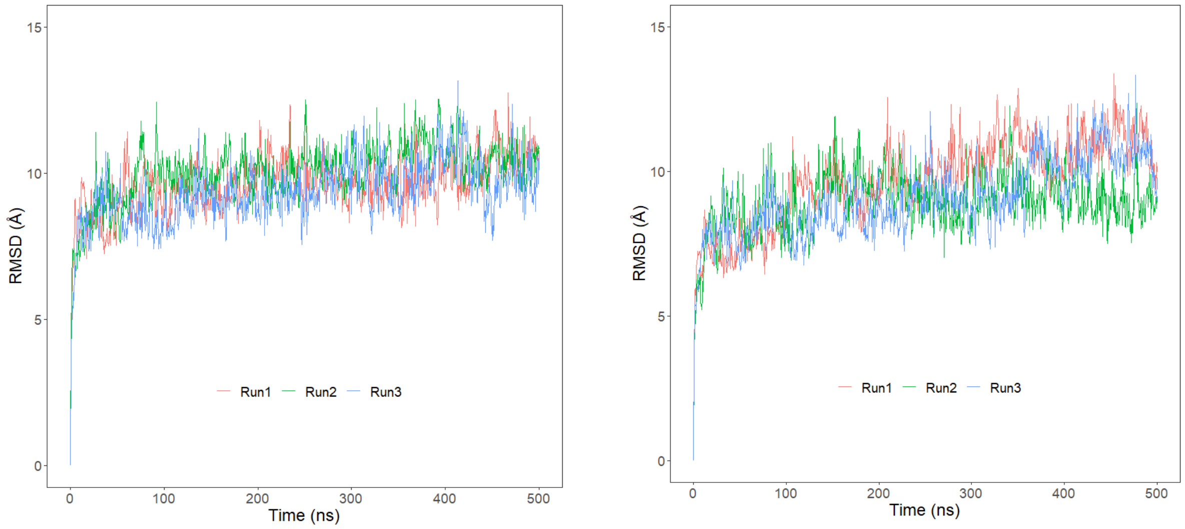


**Figure S2.** Root mean square deviation (RMSD) of protein Cα atoms with respect to the initial structure obtained from three independent runs. A) RMSD of wild type IGF-1 – IGF-1R complex; B) RMSD of mutant IGF-1 – IGF-1R complex.

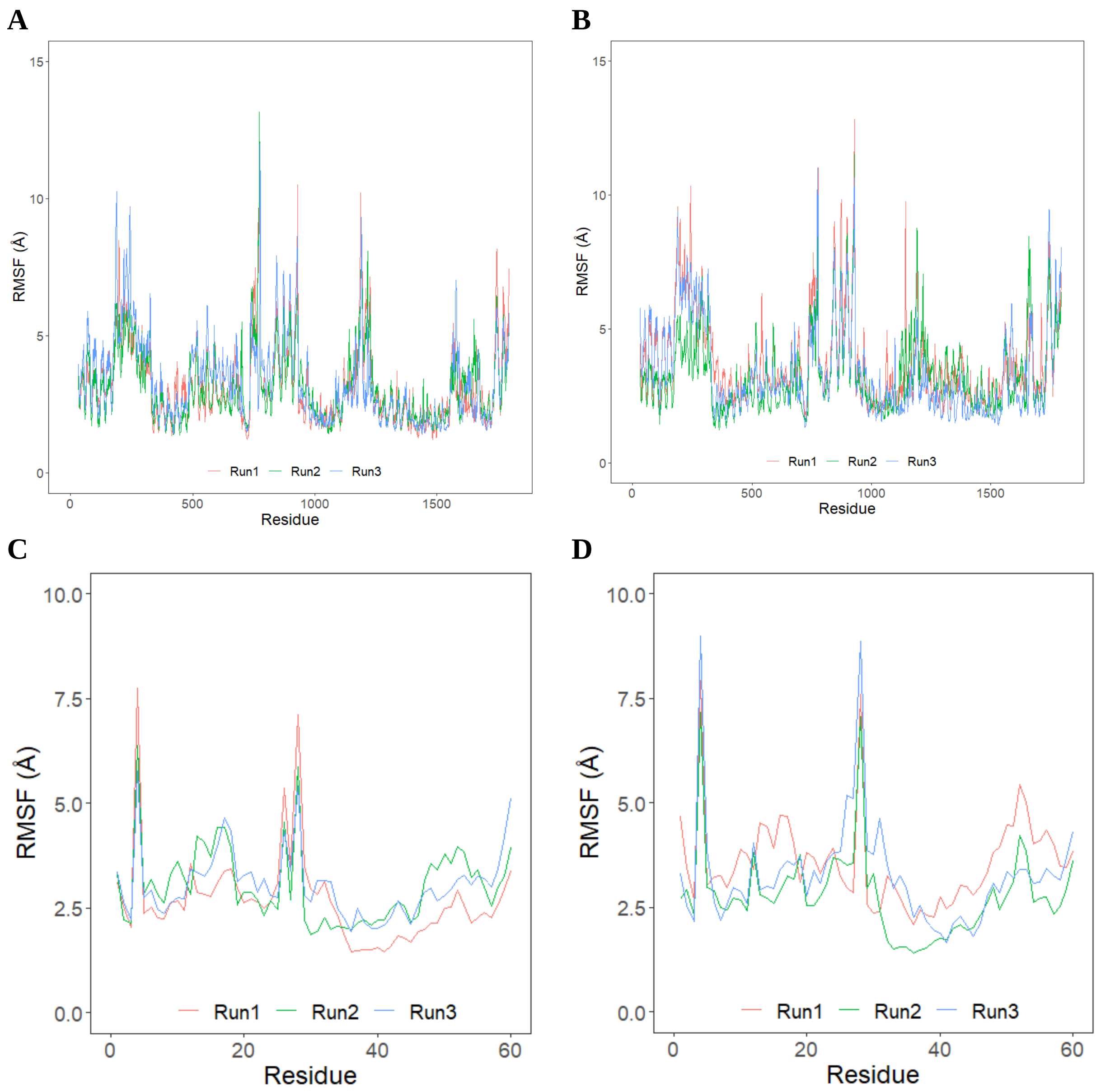


**Figure S3.** Root mean square fluctuation (RMSF) of protein Cα atoms with respect to the initial structure obtained from three independent runs. A) RMSF of wild type IGF-1 – IGF-1R complex; B) RMSF of mutant IGF-1 – IGF-1R complex; C) RMSF of wild type IGF-1; D) RMSF of mutant IGF1.

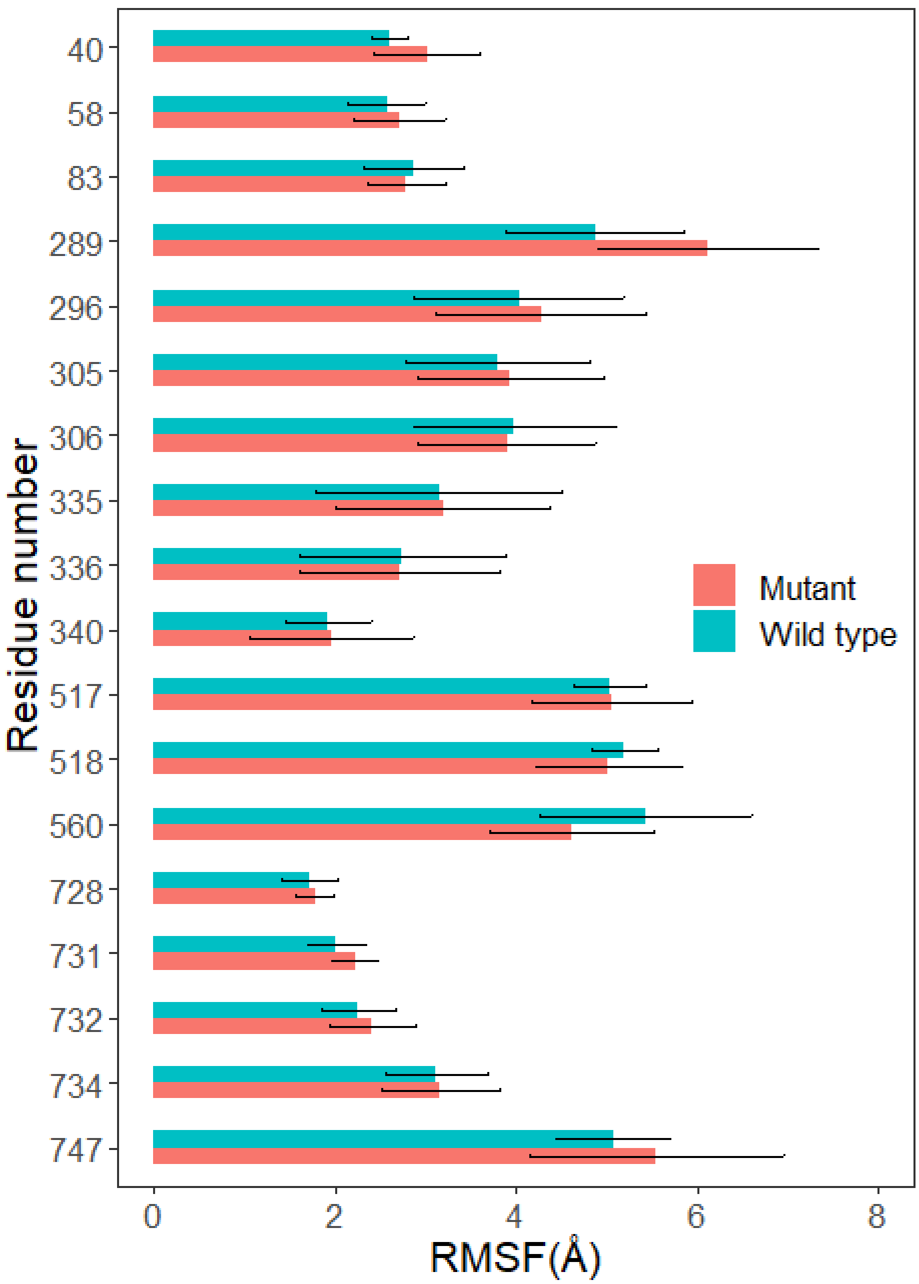


**Figure S4.** Root mean square fluctuation (RMSF) of interacting residues of IGF-1R. Results from three simulation runs of each system are plotted as mean ± SD.

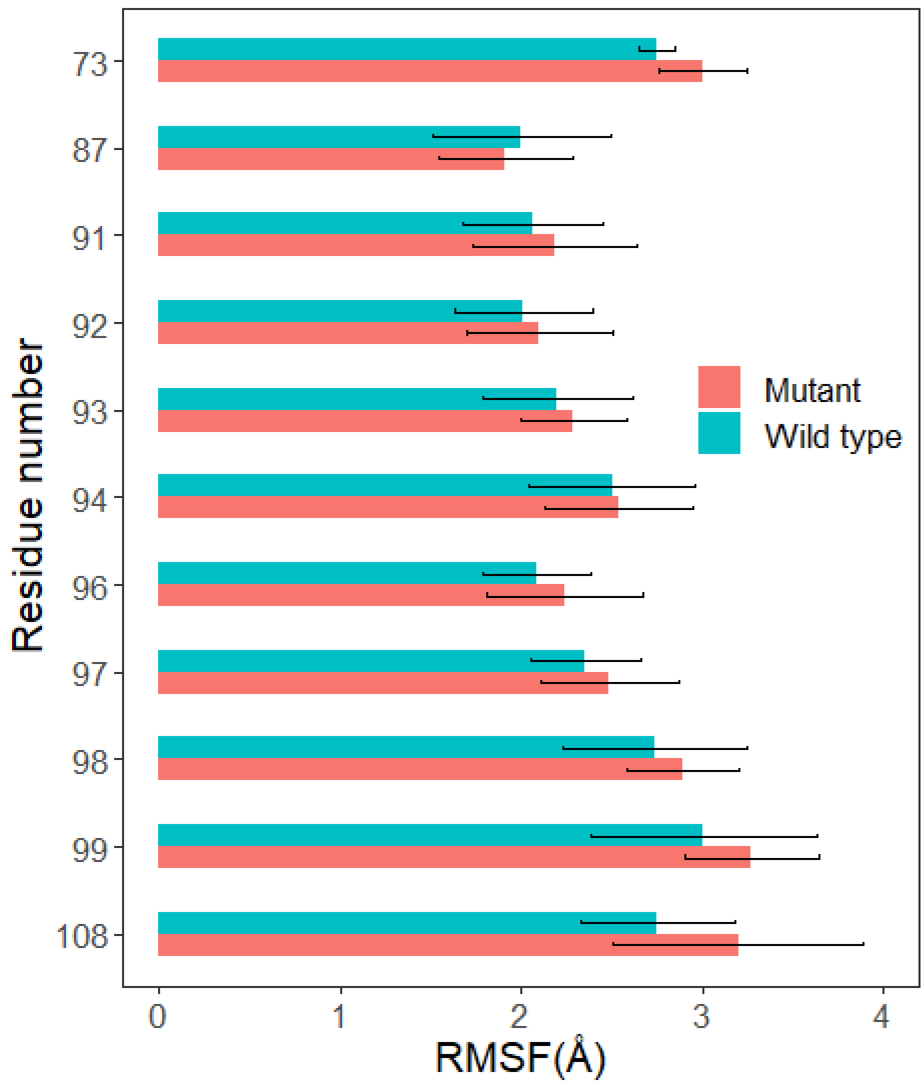


**Figure S5.** Root mean square fluctuation (RMSF) of interacting residues of IGF-1. Results from three simulation runs of each system are plotted as mean ± SD.

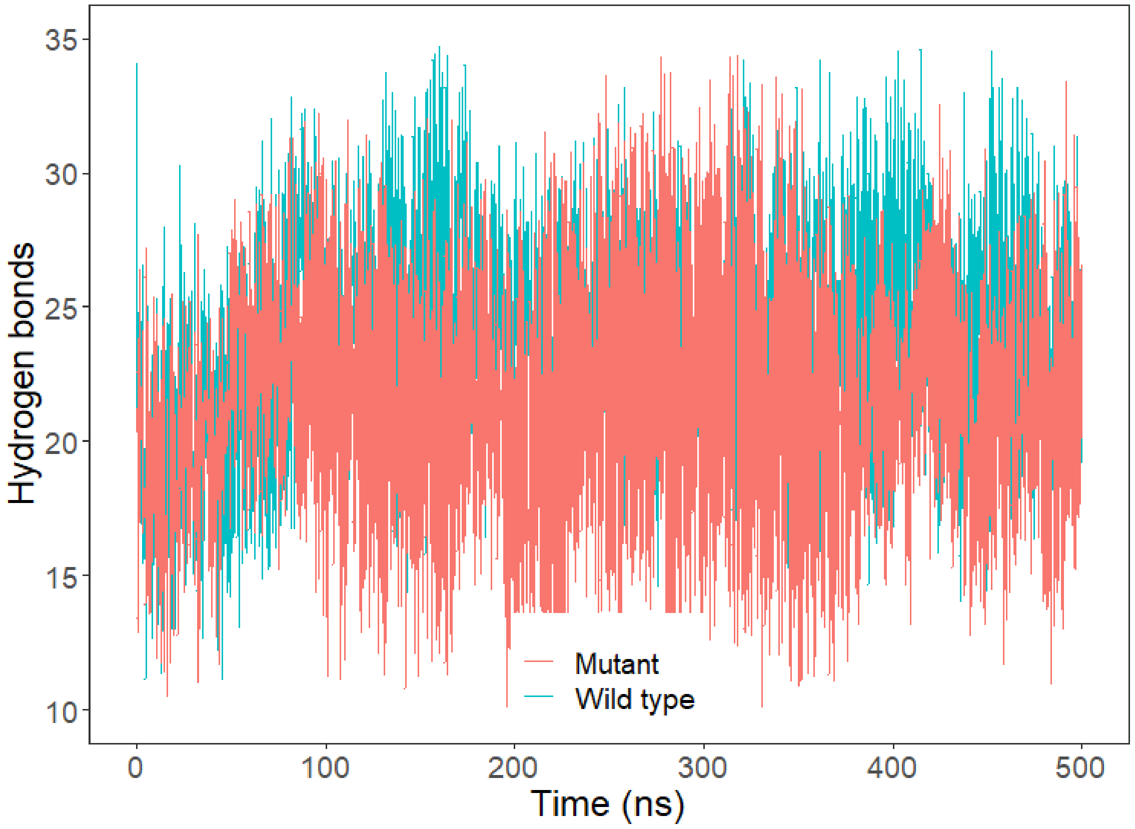


**Figure S6.** Dynamics of hydrogen bonds formation between IGF-1 and IGF-1R during simulation runs. Results from three simulation runs of each system are plotted as mean ± SD.

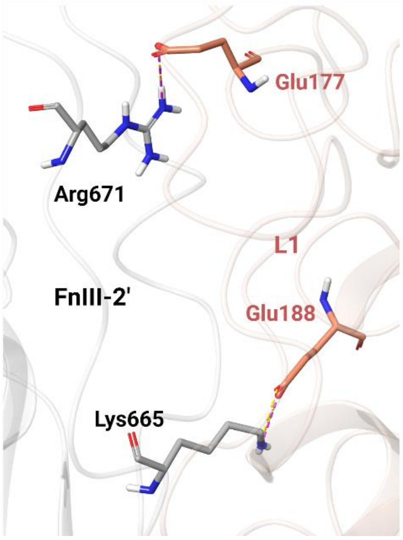


**Figure S7.** Enlarged binding pose of L1 and FnIII-2′ domains of IGF-1R. Hydrogen bonds and salt bridges are represented with yellow and pink dotted lines, respectively.

**Table S1. Cohort Characteristics.**

| Characteristics | Centenarian | Offspring | Controls |
|---|---|---|---|
| Number of participants | 686 | 1027 | 774 |
| Baseline age, mean ± SD | 97.1 ± 3.4 | 70.7 ± 7.9 | 74.8 ± 8.5 |
| Female (%) | 73.4 | 56.3 | 54.1 |
| *IGF-1:*p.Ile91Leu, (n) | 2 | - | - |
| *IGF1*:p.Ala118Thr, (n) | 2 | 4 | 3 |

**Table S2.** Radius of gyration of IGF-1R during simulation runs.

| Run | Wild type | | | | Mutant | | | |
|---|---|---|---|---|---|---|---|---|
| | Minimum | Maximum | Mean | SD | Minimum | Maximum | Mean | SD |
| 1 | 45.64 | 49.24 | 47.47 | 0.60 | 46.31 | 60.12 | 47.97 | 0.96 |
| 2 | 46.43 | 61.71 | 48.68 | 2.98 | 46.53 | 49.06 | 47.71 | 0.42 |
| 3 | 45.84 | 56.73 | 47.14 | 0.65 | 44.81 | 48.98 | 46.95 | 0.99 |

**Table S3**. Radius of gyration of IGF-1 during simulation runs.

| Run | Wild type | | | | Mutant | | | |
|---|---|---|---|---|---|---|---|---|
| | Minimum | Maximum | Mean | SD | Minimum | Maximum | Mean | SD |
| 1 | 12.83 | 13.84 | 13.30 | 0.16 | 12.93 | 14.12 | 13.48 | 0.20 |
| 2 | 12.97 | 13.95 | 13.42 | 0.17 | 12.42 | 13.79 | 13.20 | 0.25 |
| 3 | 13.19 | 14.01 | 13.60 | 0.16 | 12.83 | 13.61 | 13.23 | 0.12 |

**Table S4.** The percentage of simulation time during which intermolecular contacts were retained between IGF-1 and IGF-1R interacting residues.

| Interaction | Wild type (contact duration %) | | | | | Mutant (contact duration %) | | | | |
|---|---|---|---|---|---|---|---|---|---|---|
| | Run1 | Run2 | Run3 | Mean | SEM | Run1 | Run2 | Run3 | Mean | SEM |
| **IGF-1R:N747 – IGF-1:S99** | 95.3 | 4.5 | 68.2 | 56.0 | 21.9 | 5.0 | 8.1 | 9.3 | 7.4 | 0.9 |
| **IGF-1R:R734 – IGF-1:E94** | 91.9 | 97.8 | 56.3 | 82.0 | 10.6 | 77.7 | 31.3 | 5.0 | 37.9 | 17.4 |
| **IGF-1R:V732 – IGF-1:Y108** | 62.8 | 90.6 | 42.2 | 65.2 | 11.4 | 73.3 | 62.8 | 10.1 | 48.7 | 16.0 |
| **IGF-1R:F731 – IGF-1:I/L91** | 71.0 | 95.1 | 71.4 | 79.2 | 6.5 | 84.4 | 61.2 | 17.9 | 54.5 | 15.9 |
| **IGF-1R:N728 – IGF-1:R98** | 10.0 | 94.9 | 10.1 | 38.3 | 23.1 | 10.0 | 99.4 | 83.2 | 64.2 | 22.4 |
| **IGF-1R:N728 – IGF-1:V92** | 83.2 | 46.9 | 87.7 | 72.5 | 10.5 | 4.5 | 3.0 | 27.5 | 11.7 | 6.5 |
| **IGF-1R:N728 – IGF-1:P87** | 93.0 | 88.6 | 60.7 | 80.8 | 8.3 | 80.5 | 28.9 | 46.3 | 51.8 | 12.3 |
| **IGF-1R:K560 – IGF-1:E57** | 15.1 | 10.2 | 33.1 | 19.4 | 5.7 | 35.6 | 72.3 | 57.1 | 54.9 | 8.7 |
| **IGF-1R:R518 – IGF-1:F97** | 63.8 | 80.7 | 8.1 | 50.1 | 17.9 | 10.4 | 7.1 | 5.0 | 7.5 | 1.3 |
| **IGF-1R:Y517 – IGF-1:C96** | 94.7 | 87.9 | 89.1 | 90.6 | 1.7 | 8.1 | 4.9 | 94.1 | 35.7 | 23.8 |
| **IGF-1R:T340 – IGF-1:Q88** | 100 | 46.1 | 15.8 | 53.9 | 20.1 | 4.5 | 52.1 | 10.1 | 22.3 | 12.3 |
| **IGF-1R:K336 – IGF-1:D93** | 81.2 | 77.9 | 58.1 | 72.4 | 5.9 | 3.4 | 30.5 | 31.5 | 21.9 | 7.4 |
| **IGF-1R:E335 – IGF-1:R98** | 10.1 | 97.3 | 22.3 | 43.2 | 22.3 | 8.1 | 82.5 | 5.1 | 31.9 | 20.7 |
| **IGF-1R:E306 – IGF-1:S99** | 16.8 | 10.1 | 100 | 42.2 | 23.6 | 17.6 | 3 | 48.6 | 23.1 | 10.9 |
| **IGF-1R:Q305 – IGF-1:Y79** | 61.0 | 10.1 | 34.2 | 35.1 | 12.0 | 4.4 | 28.4 | 56.9 | 29.9 | 12.4 |
| **IGF-1R:F296 – IGF-1:Y79** | 10 | 5.1 | 14.5 | 9.8 | 2.2 | 37.1 | 33.8 | 35.8 | 35.6 | 0.76 |
| **IGF-1R:E289 – IGF-1:K75** | 31.9 | 6.5 | 10.1 | 16.1 | 6.5 | 35.7 | 47.3 | 55.5 | 46.2 | 4.7 |
| **IGF-1R:E83 – IGF-1:S81** | 63.7 | 100 | 10 | 57.9 | 21.3 | 79.1 | 10.1 | 4.9 | 31.4 | 19.5 |
| **IGF-1R:Y58 – IGF-1:G80** | 25.7 | 9.1 | 5 | 13.3 | 5.2 | 85 | 25.3 | 4.9 | 38.4 | 19.6 |
| **IGF-1R:R40 – IGF-1:F73** | 56.6 | 46.9 | 28.8 | 44.1 | 6.6 | 12.5 | 4.3 | 35 | 17.3 | 7.5 |

**Table S5.** The percentage of simulation time during which intermolecular contacts were retained between chain A and chain B interacting residues of IGF-1R dimer.

| Interaction | Wild type (contact duration %) | | | | | Mutant (contact duration %) | | | | |
|---|---|---|---|---|---|---|---|---|---|---|
| | Run1 | Run2 | Run3 | Mean | SEM | Run1 | Run2 | Run3 | Mean | SEM |
| **A:R755 - B:E563** | 36.6 | 52.2 | 2 | 30.3 | 12.1 | 5.5 | 0.04 | 94.8 | 33.4 | 25.1 |
| **A:N724 - B:R518** | 92.6 | 56.8 | 0.14 | 49.8 | 21.9 | 3 | 90.6 | 36.6 | 43.4 | 20.8 |
| **A:K720 - B:Y517** | 72.4 | 100 | 30.3 | 67.6 | 16.5 | 100 | 60.5 | 6.1 | 55.5 | 22.21 |
| **A:K709 - B:D553** | 91.4 | 60.1 | 96.3 | 82.6 | 9.3 | 57.2 | 86.9 | 48.0 | 64.0 | 9.6 |
| **A:E706 - B:D542** | 62.2 | 63.3 | 0.29 | 41.9 | 16.9 | 0 | 0 | 0 | 0 | 0 |
| **A:E690 - B:R753** | 49.7 | 0.74 | 70.6 | 40.3 | 16.9 | 12.5 | 0.29 | 0 | 4.3 | 3.4 |
| **A:K688 - B:E715** | 49.8 | 75.7 | 66.9 | 64.2 | 6.2 | 52.9 | 0.29 | 0 | 17.7 | 14.3 |
| **A:R671 - B:E177** | 7.9 | 16.8 | 93.0 | 39.3 | 22.0 | 12.6 | 4.3 | 86.0 | 34.3 | 21.2 |
| **A:K665 - B:E188** | 78.3 | 82.7 | 66.1 | 75.7 | 4.0 | 70.7 | 79.8 | 87.7 | 79.4 | 4.0 |
| **A:D616 - B:R739** | 98.1 | 98.6 | 73.3 | 89.8 | 6.8 | 45.3 | 2.7 | 72.6 | 40.2 | 16.6 |
| **A:D553 - B:R365** | 36.9 | 14.5 | 45.5 | 32.3 | 7.5 | 59.4 | 39.2 | 6.5 | 35.1 | 12.6 |
| **A:N547 - B:G367** | 11.7 | 74.9 | 0.04 | 28.9 | 23.3 | 54.1 | 3.54 | 0.09 | 19.2 | 14.2 |
| **A:R480 - B:D424** | 90.1 | 82.2 | 74.1 | 82.1 | 3.8 | 96.4 | 0.59 | 35.3 | 44.1 | 22.8 |
| **A:R391 - B:E590** | 95.4 | 33.4 | 0.2 | 43.0 | 22.7 | 7.3 | 80.5 | 87.3 | 58.4 | 20.9 |
| **A:K389 - B:E590** | 64.9 | 57.8 | 3.9 | 42.2 | 15.7 | 5 | 87.8 | 82.4 | 58.3 | 21.8 |
